# Reduced interneuron gene expression in the frontal pole cortex of schizophrenic and bipolar subjects depends on diagnosis, cell-type, and cortical layer

**DOI:** 10.64898/2026.11.23.768599

**Authors:** David M. Krolewski, Vivek Kumar, Huzefa Khalil, Matthew J. Foltz, Marquis P. Vawter, Blynn G. Bunney, Richard M. Myers, Francis S.Y. Lee, Alan F. Schatzberg, William E. Bunney, Huda Akil, Stanley J. Watson

## Abstract

Schizophrenia (SZ) and bipolar disorder (BP) represent two different psychiatric disorders with overlapping neurocognitive deficits. Postmortem studies in the prefrontal cortex indicate that GABA-ergic interneuron dysfunction may play a role in mediating cognitive symptoms in both illnesses. The lateral frontal pole cortex (LFPC) is a highly evolved prefrontal region involved in decision-making where examination of interneuron-related gene expression in SZ and BP has not been thoroughly studied. To address this gap, we performed two separate studies analyzing cortical layer-specific fluorescent *in situ* hybridization for interneuron subpopulations expressing somatostatin (SST) and reelin or cholecystokinin (CCK) and vasoactive intestinal peptide (VIP) in LFPC of SZ and BP subjects. In SZ, we show mRNA decreases were confined to upper cortical layers (SST+, CCK+ and CCK/VIP+ cells) while those for BP were additionally found in lower layers (SST+, reelin+, SST/reelin+, and CCK/VIP+ cells). This indicates that SZ and BP exhibit divergent underlying causes for the dysregulation of long-range corticocortical neurocognitive networks, corticostriatal and transthalamic circuits. However, both disorders shared a common reduction in SST+ and CCK/VIP+ cells in layer 2, indicating an overlapping system for dysregulation of long-range corticocortical networks that may affect decision-making. Overall, gene-specific mRNA decreases were more numerous in BP than those in SZ, suggesting that dysfunctional upper cortical layer-mediation of interconnected cortical network hubs may be more related to severity of neurocognitive deficits rather than total number of gene expression changes. These data suggest that LFPC interneurons in SZ and BP differentially regulate corticocortical and subcortical circuits that mediate decision-making processes.

## Introduction

The human frontopolar cortex (FPC), Brodmann Area 10, represents highly evolved cortical region which is characterized by expansive pyramidal cell spines, lower cell density, and increased columnar spacing compared to non-human primates, potentially allowing for greater connectivity with long-range corticocortical networks and subcortical-cortical circuits ^1,2^. This is particularly relevant to the lateral FPC (LFPC) where neuroimaging studies demonstrate divergent long-range functional connectivity patterns compared to monkey brain; indicating its evolutionary advancement from non-human primates ^3,4^. As such, investigation of LFPC function and its dysregulation is of special interest to better understand neuronal mechanisms associated with human neurocognition and psychiatric illness.

LFPC acts as a key hub within the frontoparietal neurocognitive network (FPN), also generally referred to as the central executive network ^5^. The FPN facilitates goal-directed behaviors through flexible interaction with the default mode (DMN) and salience (SAN) networks which are anticorrelated; engaging during interoceptive thought and cognitive tasks using externally directed attention, respectively ^6–10^. Within the FPN, the LFPC hub regulates the higher-order process of cognitive “branching” or “subgoal processing” that integrates working memory and attentional resources ^11,12^. This allows for fluid decision-making by maintaining information related to a main task in working memory while completing a secondary or alternative task ^13,14^. Specifically, while medial FPC associates more with appraisal and monitoring of the current goal, LFPC evaluates alternative task values represented by prospective episodic memory-guided scenarios that may favor more valuable options ^15,16^.

Mounting evidence suggests that the cognitive dysfunction observed in schizophrenia (SZ) and bipolar patient groups (BP) can be partly attributed to aberrant functional connectivity between nodes of the FPN, DMN and SAN-the triple network of psychopathology ^17–19^, as well as thalamocortical dysconnectivity with the LFPC ^20–23^. In support, clinical patients with LFPC lesions ^24^ experience deficits in decision-making similar to SZ and BP subjects ^25–27^. In addition, these two illnesses also display altered LFPC activity during working memory tasks ^28–32^.

Previous gene expression analyses performed within key nodes of neurocognitive networks provide a wealth of data suggesting that compromised function of specific inhibitory gamma-aminobutyric acid (GABA)-ergic cortical interneuron subtypes may underlie dysregulated local and, subsequently, long-range network communications in SZ and BP ^33–40^. While much of this work has centered around the anterior cingulate and dorsolateral prefrontal cortex (DLPFC), there has been little focus on the LPFC. Since interneuron subclasses can exhibit differential transcriptional profiles, cortical-layer positioning, and synaptic arrangements with pyramidal neurons ^41^, identifying the layer-specific dysfunction of distinct LFPC GABA-ergic cell-types is essential for better defining the circuit mechanisms underlying psychiatric symptoms.

Meta-analyses previously performed in our laboratory, which included assessment of grossly dissected LFPC tissue from SZ and BP subjects, demonstrated alterations for mRNAs corresponding to somatostatin (SST) neuropeptide-expressing interneurons as well as GABA-related proteins in SZ and BP, respectively ^42^. These findings agree with a similar non-layer-specific FPC in-situ hybridization study showing a trend for downregulation of SST mRNA in SZ ^43^.

In the current investigation, we aim to expand on previous FPC gene expression studies by: (1) performing fluorescent in-situ hybridization (FISH) specifically in LFPC region of both SZ and BP subjects for SST and additional interneuron subtype markers including the glycoprotein reelin, cholecystokinin (CCK) peptide, and vasoactive intestinal peptide (VIP) and, (2) quantifying cortical layer-specific expression of each gene and their colocalization across diagnoses. These results will further refine our understanding of potential differences in LFPC interneuron-mediated circuitry associated with neurocognitive deficits in two psychiatric disorders.

## Methods

### Diagnostic procedures

A full psychological autopsy on all subjects was performed postmortem according to previous literature ^44^ with SZ and BP diagnoses meeting requirements of the Diagnostics Manual of Mental Disorders IV ^45^. Evaluation of medical examiner/coroner records, psychiatric records, toxicology reports, history of substance abuse, physical health, and next-of-kin interviews to obtain additional details regarding symptomology were considered. Control subjects or first-degree relatives did not present evidence of neurological or psychiatric disorders.

### Brain dissection

General dissection methods have been previously described ^46^. Briefly, tissue was obtained from the Brain Donor Program at the University of California, Irvine.

Following brain removal at autopsy, coronal slabs of ∼0.8 cm were cut, flash-frozen, and stored at −80°C. Blocks were sectioned at 30 µm in a cryostat, thaw mounted onto Superfrost^TM^ Plus glass slides (Fisher Scientific, Pittsburgh, PA) and stored at −80 °C until utilized. All subjects had zero agonal factors with postmortem interval (PMI) of <37 h and, except for one subject with unknown pH, all subjects had pH above 6.5 to maximize mRNA preservation ^47,48^.

### Fluorescence in situ hybridization chain reaction (HCR-FISH)

Two separate in situ experiments (ISH) were conducted. DNA probes for SST and reelin were simultaneously implemented in the first assay and those for CCK and VIP were combined in the second experiment. Split initiator DNA probes for ISH were designed in our lab and synthesized by Integrated DNA Technologies (Coralville, Iowa)^49^. Full details of probe design, hybridization, amplification, and imaging conditions are provided in Supplementary Methods.

### Statistical Analysis

For gene expression analysis in the LFPC of control subjects only, a Mann-Whitney-U Test between layers was performed with p-values FDR corrected. To assess whether a diagnosis of SZ or BP resulted in gene expression changes, a mixed model analysis with negative binomial regression was conducted by letting the intercept vary by subject (random effect on subject). A separate model was used for each gene. The adjusted count density was the dependent variable while the diagnosis and layer and their interaction were the independent variables. Any data points which were ±3 standard deviations away from the mean were considered outliers and removed. After correlation analysis it was determined that age should be added for CCK and VIP as a covariate and freezer storage time be added as a covariate for all the genes analyzed. The mixed model analysis was conducted using the glmmTMB package ^50^ in R and the between diagnosis and layer contrasts were calculated using the emmeans package ^51^. The p-values are corrected using the Tukey method across the contrasts. The degrees of freedom are estimated using Satterthwaite’s method using the package lmerTest ^52^. The p-values are corrected using the Tukey method. All results were considered statistically significant at p < 0.05. Post-hoc data points ±3x the standard deviation were considered outliers and removed.

### Imaging and quantification

Image z-stacks of 30µM-thick brain sections were acquired using an Olympus Fluoview-3000 confocal microscope. Quantitative analysis of captured ISH images demonstrating positive signal in defined cortical layers was accomplished with using a custom Image J filter designed to segment and filter labeled cells. See Supplemental Methods for detailed protocol.

## Results

### Covariate results from negative binomial mixed model

An effect of cortical layer was found for all genes: SST (χ^2^=537.628, Df=2, N=55 p<0.001), reelin (χ^2^=2171.399, Df=6, N=54, p<0.001), CCK (χ^2^=826.601, Df=6, N=54, p<0.001), and VIP (χ^2^=1745.762, Df=6, N=56, p<0.001). For individual genes, an overall effect of diagnosis on gene expression was only significant for SST χ^2^=6.210, Df=2,N=53, p=0.045). However, reelin expression exhibited a significant diagnosis/layer interaction (χ^2^=22.466, Df=12, N=54, p=0.033). VIP showed an effect of age (χ^2^=11.207, Df=1, N=56, p=0.001). For colocalization density of double labeled neurons, SST/reelin cells, exhibited an effect of layer (χ^2^=741.344, Df=6, N=52, p<0.001) and a diagnosis/layer interaction (χ^2^=24.901, Df=12, N=52, p=0.015). Colocalized CCK/VIP cells exhibited an effect layer (χ^2^=2036.586, Df=6, N=56, p<0.001) and diagnosis (χ^2^=8.801, Df=2, N=56, p=0.012).

### Control Subjects: Visualization of gene expression and relative abundance by cortical layer

Following FDR correction, the Mann-Whitney-U Test in control subjects showed cortical layer differences in cell count density based on gene/layer interaction (**Supplemental Fig. 1A-D**). The density of cells expressing SST in cortical layer 2 was higher than all other layers except layer 4 which had the next highest count. Layer 6b exhibited the lowest median cell count which was significantly lower compared to all other layers. The density of reelin-positive cells in cortical layer 1 was larger than that of all other individual layers. The density of reelin-expressing layer 4 cells was higher compared to layer 3 and lower layers 5, 6a, and 6b. CCK-positive cells in cortical layers 2, 3, 4, and 6a were not different from each other but significantly more abundant compared to layers 1, 5, and 6b. Cells expressing VIP were significantly higher in layer 2 compared to all other layers. VIP-positive cells positioned in layers 1, 3, and 4 were more densely expressed than in layers 5, 6a, and 6b

Further quantitative analysis was performed for total density of double-labeled cells across layers exhibiting colocalization of SST/reelin or CCK/VIP (**Supplemental Fig. 2A,B**. The cellular density of SST colocalization with reelin in cortical layers 1 and 4 was significantly higher than all other cortical layers, having cell counts greater than remaining layers 2, 3, 5, 6a, and 6b.

Lastly, the median percent colocalization as a percentage of total cells expressing each gene was analyzed in control subjects (**Supplemental Fig. 3A-D)**. The percentage of SST-expressing cells colocalizing reelin was significantly higher in layers 1 and 4 versus all other layers at 74% and 57%, respectively. For the reverse combination, reelin/SST, the median number of positive cells was highest in layers 4 and 5 at 67% and 71%, respectively. The percentage of CCK-positive cells exhibiting co-expression of VIP were highest in layers 1 and 2 (18% and 15%, respectively). VIP-positive cells co-expressing CCK showed a gradient of colocalization .The greatest percent colocalization was in layers 1 and 2 (43% and 44%, respectively) demonstrating significant differences versus layers 4, 5, 6a, and 6b. Layer 3 colocalization was intermediate at 34% and was also significantly higher than layers 5, 6a, and 6b.

### Effect of Diagnosis: Pairwise post-hoc comparisons using negative binomial mixed model ANOVAs

When SST gene expression was quantified with all layers combined, SST showed a significant reduction in BP (p=0.037), data not shown. SST gene expression was more specifically downregulated in layer 1 of both SZ (p=0.028, **Fig. 3A**) and BP subjects relative to controls (p=0.015, **Fig. 3A**) with additional significance in layers 4 (p=0.023, **Fig. 3B**) and 5 (p=0.033, **Fig. 3C**) for the BP group.

Reelin expression was lower in BP versus SZ patients in layer 5 (p=0.035, **Fig. 3D**) and layer 6b (p=0.038, **Fig. 3E**).

CCK expression was only significantly downregulated in layer 2 of SZ subjects relative to controls (p=0.017, **Fig. 3F**).

VIP expression showed no significant differences in either SZ or BP subjects relative to control subjects.

Further quantitative analysis of double-labeled cells across layers assessed the impact of diagnosis on both colocalization SST/reelin or CCK/VIP neurons. In BP subjects, when all layers were combined SST/reelin cells were significantly fewer in the BP group (p=0.044, data not shown). More specifically SST/reelin cells were diminished in layers 1 (p=0.047, **Fig. 4A**), 4 (p=0.014, **Fig. 4B**), and 5 (p=0.029, **Fig. 4C**). Also in BP subjects, combined layer analysis showed a significant reduction CCK/VIP double-label cells (p=0.011, data not shown) and were reduced in layers 1 (p=0.005, **Fig. 4D**), 2 (p=0.024, **Fig. 4E**), and 6a (p=0.017, **Fig. 4F**). In SZ subjects, the effect was more selective, as decreased CCK/VIP cell density was confined to layer 2 (p=0.030, **Fig 2E**). A summary schematic of gene expression results and their relationship to LFPC circuitry is shown in **Fig. 5**.

**Fig. 1.**
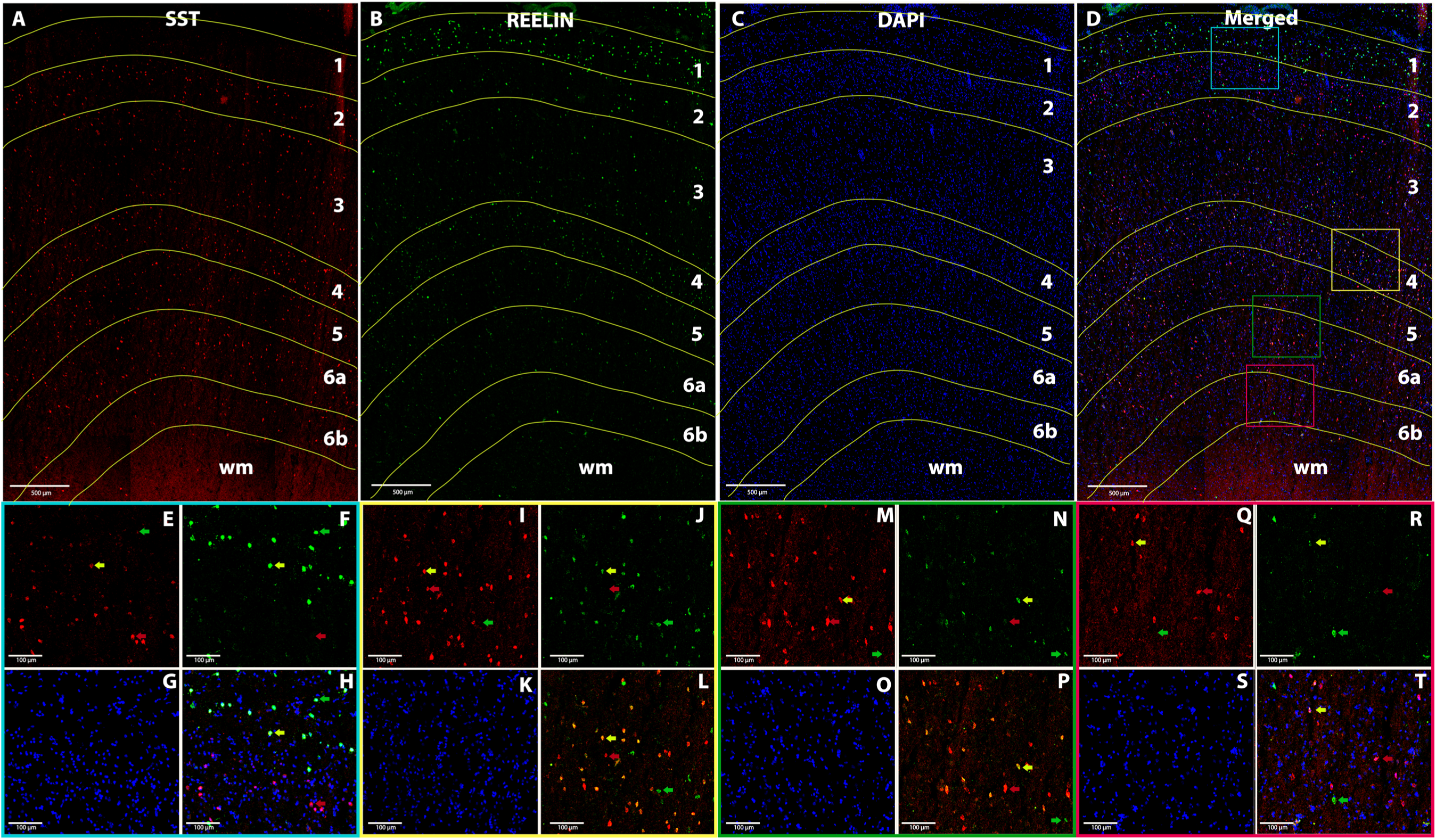
Visualization of cortical layer-based gene expression for SST and reelin in the LPFC. **A**. Individual layer expression of SST (red), **B.** Reelin (green), **C.** Nucleus marker dapi (blue), and **D.** Color-coded boxes chosen for the layer-based higher magnification pictures of the three merged signals depicted in the bottom two rows. High magnification visualization of **E.** SST, **F.** reelin, and **G.** dapi with **H.** showing all signals merged in layers 1/2. Red arrows point to single labeled SST cells, green arrows represent single labeled reelin cells, and yellow arrows demonstrate colocalization of SST and reelin. High magnification visualization of **I.** SST, **J.** reelin, and **K.** dapi, in layer 4 with **L.** showing all signals merged. High magnification visualization of **M.** SST, **N.** reelin, and **O.** dapi, in layer 6a with **P.** showing all signals merged. High magnification visualization of **Q.** SST, **R.** reelin, and **S.** dapi, in layer 6b with **T.** showing all signals merged. Dapi; 4’6-diamidino-2-phenylindole, LFPC; lateral frontal pole cortex, SST; somatostatin, wm; white matter.

**Fig. 2.**
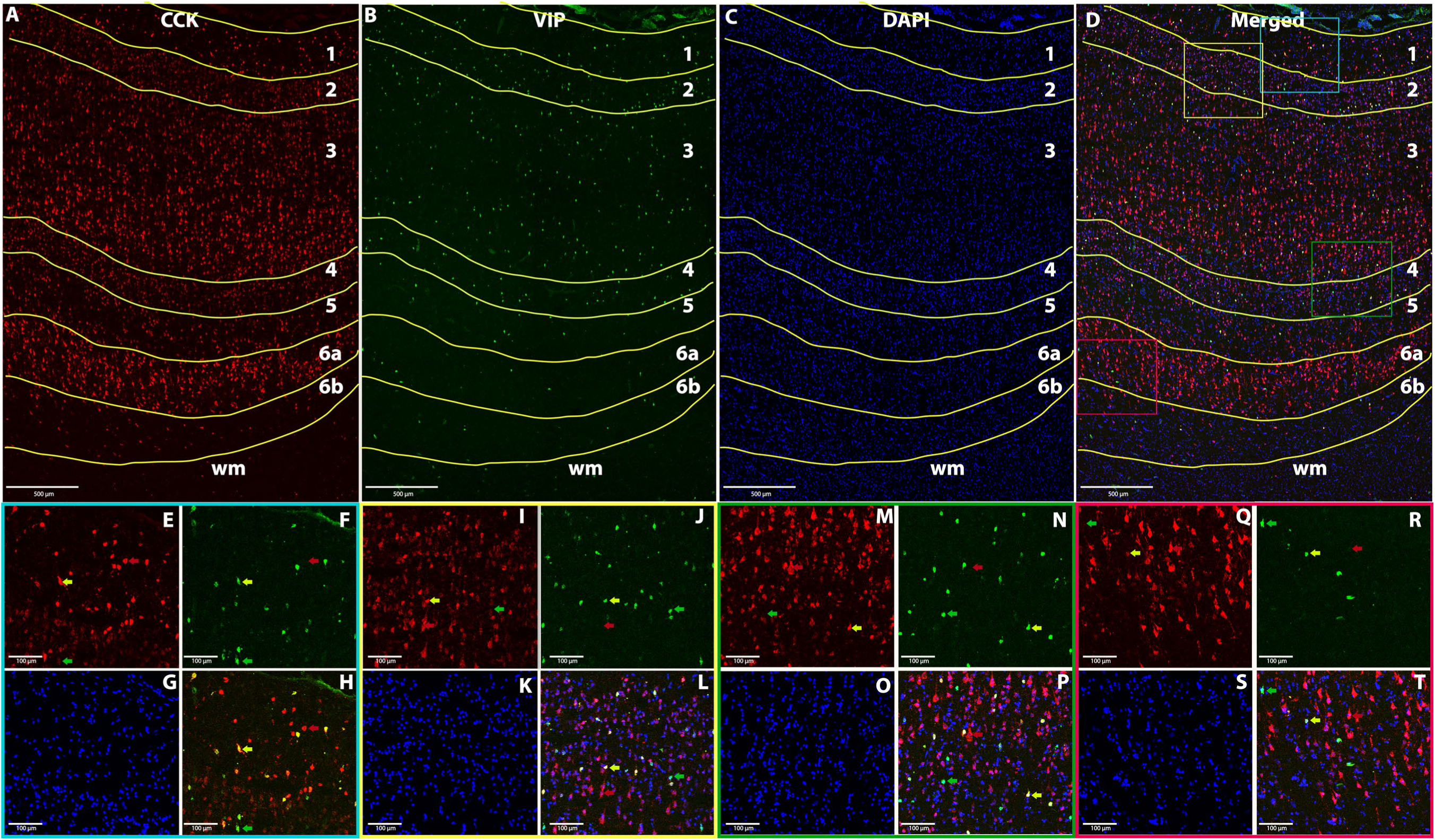
Visualization of cortical layer-based gene expression for CCK and VIP in the LPFC. **A**. Individual layer expression of CCK (red), **B.** VIP (green), **C.** Nucleus marker dapi (blue), and **D.** Color-coded boxes chosen for the layer-based higher magnification pictures of the three merged signals depicted in the bottom two rows. High magnification visualization of **E.** CCK, **F.** VIP, and **G.** dapi, in layer 1 with **H.** showing all signals merged. Red arrows point to single labeled CCK cells, green arrows represent single labeled VIP cells, and yellow arrows demonstrate colocalization of CCK and VIP. High magnification visualization of **I.** CCK, **J.** VIP, and **K.** dapi with **L.** showing all signals merged in layer 2/3. High magnification visualization of **M.** CCK, **N.** VIP, and **O.** dapi, in layer 3/4 with **P.** showing all signals merged. High magnification visualization of **Q.** CCK, **R.** VIP, and **S.** dapi, in layer 6a with **T.** showing all signals merged. CCK; cholecystokinin, Dapi; 4’6-diamidino-2-phenylindole, LFPC; lateral frontal pole cortex, VIP; vasoactive intestinal peptide, wm; white matter.

**Fig. 3.**
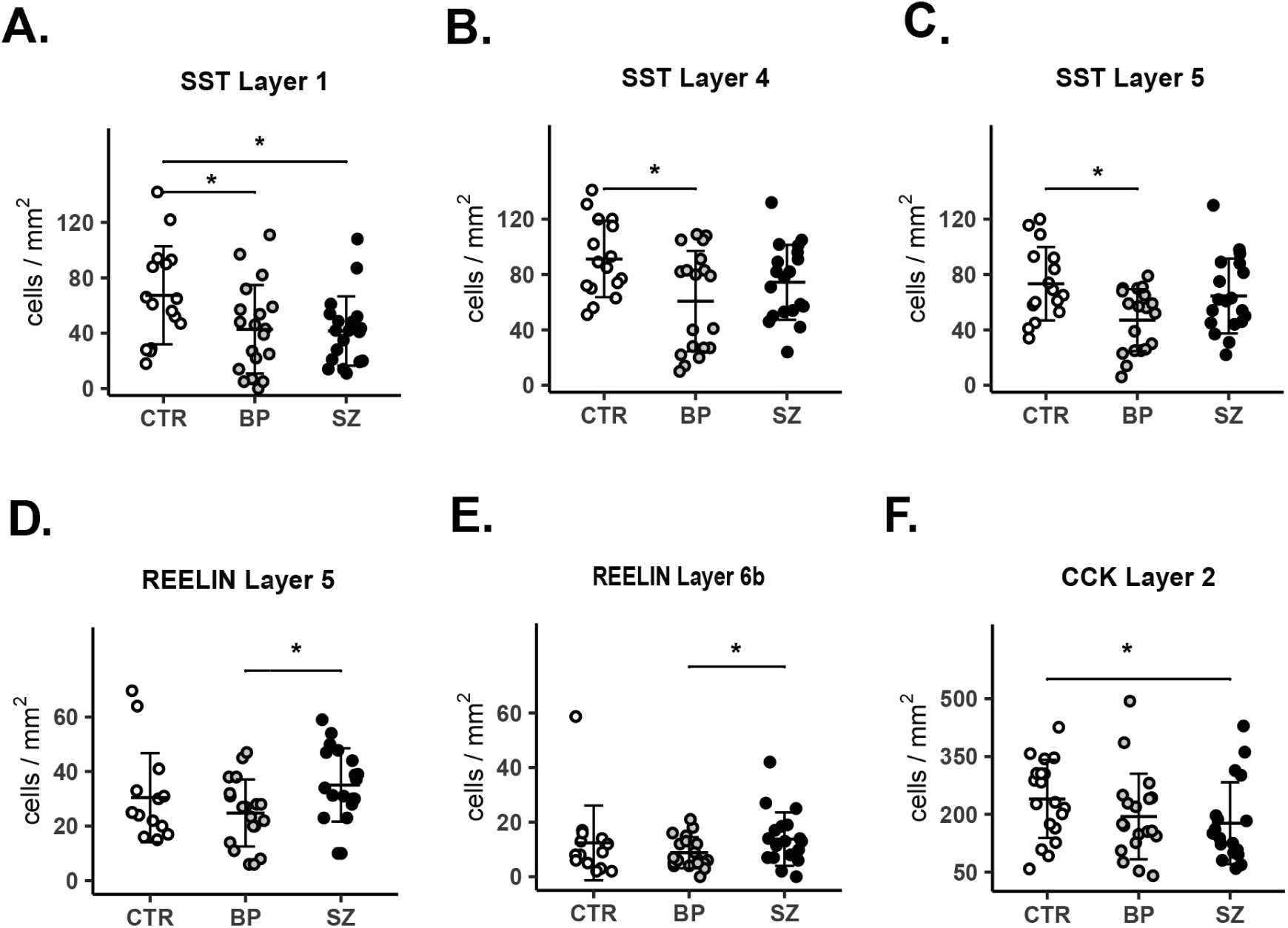
Decreased density of cells expressing either SST, reelin, or CCK mRNA as assessed by cortical layer in the LFPC of psychiatric and control subjects. **A.** SZ and BP subjects both showed reduced SST gene expression versus CTR in layer 1 with p=0.028 and p=0.015, respectively. **B, C.** BP subjects demonstrated decreased SST mRNA versus CTR layer 4, p=0.023 and layer 5, p=0.033, respectively. **D.** Reelin gene expression was reduced in layer 5 compared to SZ subjects, p= 0.035. **E.** Reelin mRNA was decreased in layer 6b versus SZ subjects, p=0.038. **F.** CCK mRNA was reduced in layer 2 of SZ subjects compared to CTR, p=0.017. Each dot represents one subject, and data is presented as mean ± standard deviation. Asterisk denotes p<0.05. BP; bipolar disorder, CCK; cholecystokinin, CTR; control, LFPC; lateral frontal pole cortex, SZ; schizophrenia, SST; somatostatin

**Fig. 4.**
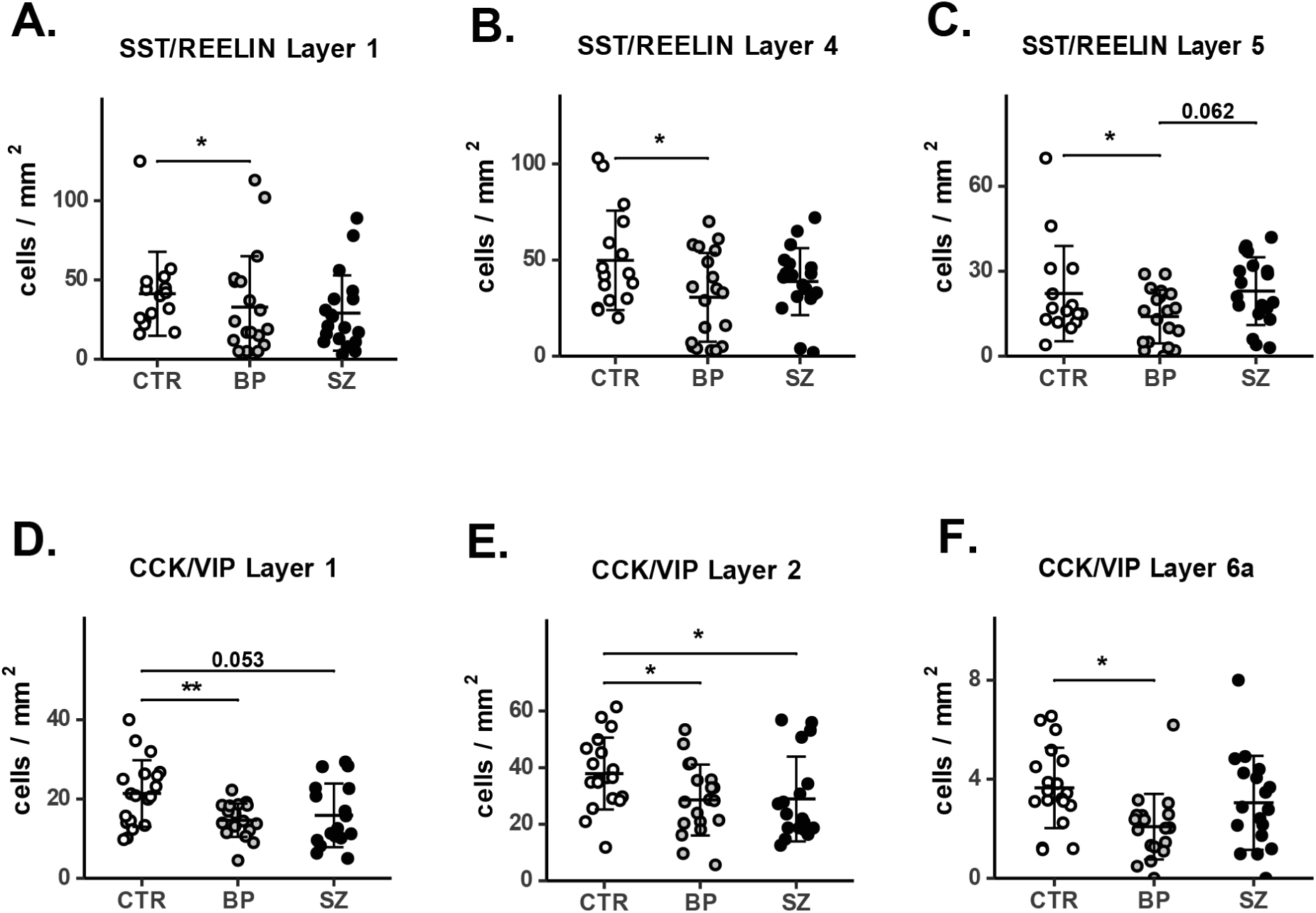
Decreased density of cells co-expressing either SST/reelin or CCK/VIP as assessed by cortical layer in the LFPC of psychiatric and control subjects. **A.** BP subjects showed reduced density of SST/reelin co-expressing cells versus CTR in layer 1, p=0.047. **B.** BP subjects demonstrated decreased SST/reelin cells versus CTR layer 4, p=0.014. **C.** SST/reelin colocalization was reduced in layer 5 versus CTR, p= 0.029 with a trend for reduction compared to SZ subjects, p=0.062. **D.** BP subjects exhibited a reduction in CCK/VIP colocalization in layer 1, p=0.005 with a trend for SZ versus CTR, p=0.053. **E.** CCK/VIP cell density was decreased in layer 2 of SZ and BP subjects, p=0.024 and p=0.030, respectively. **F.** CCK/VIP colocalized cells were decreased in layer 6a of BP subjects, p=0.017. Each dot represents one subject, and data is presented as mean ± standard deviation. Asterisk denotes p<0.05. BP; bipolar disorder, CCK; cholecystokinin, CTR; control, LFPC; lateral frontal pole cortex, SST; somatostatin, SZ; schizophrenia, VIP; vasoactive intestinal peptide.

**Fig. 5.**
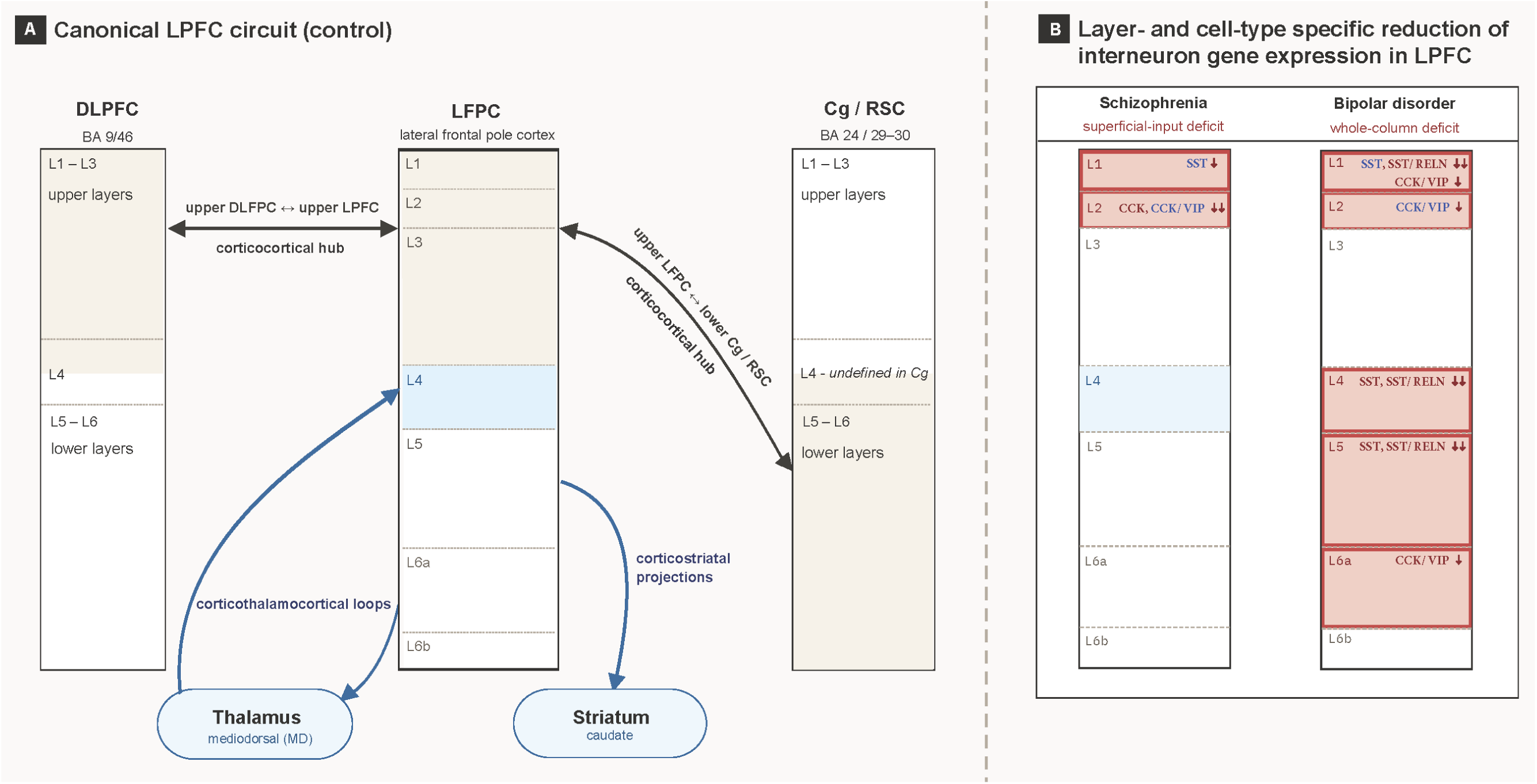
Layer-resolved LFPC circuits and differentially reduced interneuron gene expression in SZ and BP may uniquely alter reciprocal upper and lower cortical layer-mediated circuits. **A.** Highly evolved LFPC and DLPFC eulaminate cortices would be expected exhibit a higher degree of reciprocal upper layer corticocortical connections, as compared to intercommunications between upper and lower cortical layers, potentially forming a cortical hub important for mediating higher-order neurocognitive function. Alternatively, LFPC long-range cortical connections with more primitive cortices, such as the Cg/RSC, are predicted to establish reciprocal corticocortical circuits more prominently with lower cortical layers. The latter circuit arrangement may subsequently contribute to the regulation of corticothalamocortical loops and corticostriatal circuits. **B.** SZ and BP subjects both demonstrated gene expression reductions in upper cortical layers 1 and 2, having common alterations in layer 1 SST+ cells and layer 2 cells co-expressing VIP-CCK mRNA (highlighted in blue). In contrast to SZ, BP subjects displayed additional reductions in SST and reelin mRNA within layers 4 and 5 which may provide a source for the dysregulation of thalamocortical-driven circuits and corticostriatal systems, respectively. BP subjects also showed decreased VIP-CCK+ cells in layer 6a, suggesting a possible role in the dysfunction of corticothalamic-mediated transmission. BA; Brodmann area, Cg; cingulate cortex, CCK; cholecystokinin, DLPFC; dorsolateral prefrontal cortex, LFPC; lateral frontal pole cortex, MD; mediodorsal thalamus, RELN; reelin, RSC; retrosplenial cortex, SST; somatostatin, VIP; vasoactive intestinal peptide

Percentage of co-expression by cell-type was assessed by cortical layer and we found differences based on diagnosis. Reelin-positive cells expressing SST exhibited a trend for a lower degree of colocalization percentage in layer 5 of BP subjects versus CTR (p=0.076, **Supplemental Fig. 4A**) and SST cells expressing reelin showed diminished colocalization percentage in BP subjects versus SZ in layer 6b (p=0.051, **Supplemental Fig. 4B**). CCK-positive cells had a significantly lower percentage of colocalization with VIP in layer 6a of BP subjects (p=0.015, **Supplemental Fig. 4C**).

## Discussion

The present study was conducted to uncover both shared and unique features within the LFPC-mediated circuitry in SZ and BP patients, as both groups of patients exhibit symptoms of psychosis along with different degrees neurocognitive deficits, but also have distinct characteristics in mood, perception, and function ^53,54^. Both SZ and BP groups exhibited reductions in SST mRNA in cortical layer 1, and in cells colocalizing CCK/VIP in layer 2, suggesting a possible role of these layer-specific neuron types in psychosis. However, our data revealed largely non-overlapping GABA-ergic interneuron-related gene expression reductions within functionally distinct cortical laminae for each diagnosis, with the more appreciable gene expression effects being associated with BP rather than SZ, particularly in lower layers. Thus, our results point to largely diagnosis-specific dysregulation of local circuitry that may underpin differential modulation of long-range corticocortical, corticostriatal, and transthalamic communications between disorders. In the following text we offer a framework wherein reduced gene expression in specific GABA-ergic cell-types within distinct cortical laminae may impact LFPC neurocognitive deficits concerning learning, cognitive control, and decision-making in these two psychiatric illnesses.

### Somatostatin expression

In control subjects, SST-positive cell density was generally consistent across layers, except for layer 6b which had exceptionally low expression, whereas layer 2 had more expression than all other cortical layers stressing its particular importance in mediating long-range corticocortical communications (discussed below). By diagnosis, our assessment of SST gene expression revealed diminished levels in layer 1 of both SZ and BP but with greater overall cortical layer-specific deficits associated with BP in layers 4 and 5. This aligns with the lower BP group means for SST mRNA signal found in remarkably widespread neuroanatomical analyses of dorsolateral prefrontal cortex (DLPFC), orbital frontal cortex, posterior cingulate, and retrosplenial cortices relative to SZ patients ^37,40^. While strongly implicating SST in BP cortical network dysfunction, these results seemingly go against a simpler hypothesis that more severe cognitive symptoms may be more associated with greater interneuron dysfunction, considering SZ is thought to have overlapping cognitive deficits of greater scale compared to BP ^53,54^. This questions to what extent the degree of local interneuron dysfunction, across cortical regions, relates to the severity of symptoms in psychiatric illness. Yet, the present findings concur with our previous meta-analysis of grossly dissected LFPC in SZ subjects showing significantly reduced SST mRNA as well as the non-significant trend for diminished SST in SZ concluded from an additional in situ hybridization study^43^.

Most SST cortical interneurons are Martinotti cell-types that engage in fast inhibition through synapses on the dendrites of excitatory glutamatergic pyramidal cells in upper cortical layers as well as those in lower layers 5 and 6 ^41^. In human cortex, SST+ cells are represented by a variety of morphological shapes, suggesting the possibility of some degree of non-Martinotti cell inclusion in our analysis ^55^. Nevertheless, SST expressing cells are in anatomical position to modulate pyramidal cell dendrite activity that regulates long-range corticocortical and subcortical-cortical terminations upon pyramidal cells ^56,57^. This concept implies potential dysregulation of circuit synchronization through previously described influence over gamma and/or theta oscillatory activities ^57,58^ which are altered in neuropsychiatric disorders, albeit most reported in SZ patients ^59,60^.

Our findings revealing decreased SST expression in cortical layer 1 of both SZ and BP groups, along with reductions in middle/lower layers 4 and 5 exclusively in BP, highlights the importance of understanding the behavioral consequences of regional abnormalities. For example, layer 1 predominantly consists of apical dendritic tufts of layers 2, 3, and 5 pyramidal cells and axon terminals from distant cortical regions as well as those derived from subcortical nuclei producing acetylcholine, noradrenaline, dopamine, and serotonin ^61^. In addition, lamina 1 is also a primary site for integration of local amygdala afferents ^62–64^, forming a LFPC-amygdalar circuit hypothesized to gate emotional control related to decision-making ^65^.

Taken together, SZ and BP disorders may share a dysregulated sharpening of diverse and convergent signaling at the dendritic level due to compromised SST+ interneurons. For both diagnoses, this may contribute to disorganization of the proposed top-down and bottom-up integration functions of layer 1 that mediate decision-making components including task context, expectations, learning, emotion, and attention ^61,65,66^. Indeed, SZ and BP patients have both been shown to exhibit difficultly completing cognitive tasks in the presence of irrelevant emotional stimuli ^67^. It is also possible such disorganization leads to disruption of layer 1-mediated corticocortical neurotransmissions between LFPC and other nodes of long-range neurocognitive networks, hence contributing to the subsequent abnormal functional interconnectivity between the FPN and DMN/SAN that is well known to exist in both SZ and BP ^17–19^.

While cortical lamina 1 has also been described as a recipient of thalamocortical terminals ^55,61^, results from primate studies demonstrate robust thalamic afferents to highly evolved cortices, i.e., “eulaminate” type, like the LFPC, strongly innervate middle layers including layer 4 ^68–70^. In this layer, predominantly non-Martinotti SST+ interneurons synapse principally upon inhibitory parvalbumin-positive interneurons and act to disinhibit local excitatory pyramidal neurons ^41^. Therefore, the reduced SST expression we observed in layer 4 of BP subjects would suggest altered excitatory synchrony between the of LFPC pyramidal neurons and thalamocortical inputs from the mediodorsal thalamus which provides the strongest subcortical input to LFPC ^70,71^ and carries similar information related to spatial working memory, inhibitory control, and decision-making ^72,73^.

While the strongest evidence for thalamo-prefrontal cortical neuropathology exists for SZ patients ^74^, those diagnosed with SZ or BP with psychosis each display heightened symptoms positively correlated with reduced thalamocortical functional connectivity to the LFPC ^75^. Thus, the compromised LFPC layer 4 SST gene expression in BP may also, in part, play a role in the mechanisms underlying thalamocortical functional hypoconnectivity involving LFPC in BP as reported in other studies ^20,23,76^. This may also help explain, along with overlapping layer 1 SST expression changes, some of the common symptomology with SZ patients because BP subjects also show deficits in working memory, planning, and set shifting ^32,69,77^. Interestingly, the reduced SST mRNA previously shown in DLPFC layer 4 of SZ subjects ^39,78^ suggests that dysregulation of thalamocortical circuits associated with different diagnoses may lead to similar neurocognitive deficits, although perhaps mediated by different but highly interconnected neuroanatomical regions of the FPN network, i.e. LFPC and DLPFC.

Showing further consistency in BP, we also uncovered reduced SST mRNA in layer 5. This layer houses diverse pyramidal projection neurons that form corticocortical, corticostriatal, and corticothalamic circuits. As such, we suspect dysregulated SST+ may contribute to some reduced ability to tune layer 5 excitatory pyramidal neurons, perhaps synapsing on local dendrites in addition to distal dendrites that reach layer 1 through ascending branching ^79^. However, the magnitude of functional effects on long-range corticocortical networks is questionable. Following the prediction of intracortical connection patterns of primate prefrontal cortex ^80^, layer 5 corticocortical neurons in LFPC would be expected to have only modest interconnectivity with DLPFC, a network hub of the FPN, while likewise providing a minimal feedforward efferent input to the cingulate gyrus that encompasses nodes from the SAN and DMN ^6,9^. Therefore, dysregulated SST+ interneurons in layer 5 are not likely to remarkedly affect the functional connectivity of LFPC corticocortical output neurons interacting with the SAN and DMN through known LFPC connections with the anterior midcingulate and the pregenual cingulate/retrosplenial cortices, respectively ^81–84^.

On the other hand, there is evidence indicating that disrupted SST+ interneurons are more likely to alter the electrical activity of layer 5 corticostriatal neurons ^85,86^. Interestingly, BP subjects demonstrate abnormalities in prefrontal-striatal systems associated with set-shifting and impulsive decision-making ^87,88^. In line with this, in addition to the MD, layer 5 pyramidal neurons project to the ventral anterior/ventrolateral motor thalamus region ^89^.

### Reelin expression

In control subjects, reelin-positive cells were especially concentrated in layer 1 highlighting a potential specialty in gating top-down bottom-up information processing. By diagnosis, contrasting from the pattern of diminished SST gene expression in BP versus controls, we demonstrate reelin is decreased in BP compared to SZ subjects in layer 5 as well as 6b, differentiating expression by diagnosis and layer. These findings agree with one study showing reduced reelin+ immunostaining and mRNA signal in BA9 of BP subjects ^90^ but are distinct from previous reports in SZ versus controls, describing decreased reelin mRNA found in pooled cortex derived from combined BA46 and non-regionally defined FPC ^90,91^. In human cortex, reelin and SST have been previously shown to be co-expressed in GABA-ergic interneurons ^92^. Rodent studies show that most reelin+ cells exhibit considerable colocalization with SST, about 30-50%, at both the mRNA and protein levels ^93–97^.

In our analysis, we found a similar percentage of total co-expression with SST within the LPFC (all layers combined) at 37% for the mean in control subjects, in agreement with previous cortical studies in rodent. Layer assessment of reelin-positive cells co-expressing SST revealed that layers 4 and 5 had the highest percentage of colocalization at 64% and 68%, respectively. These findings indicate reelin/SST cell-types play a particular importance in shaping thalamocortical input and corticostriatal output. By contrast, the percent of SST-positive cells demonstrating colocalization with reelin was analyzed and layers 1 and 4 exhibited the highest total cell count percentage at 74% and 57%, respectively. This reinforces the relative importance of reelin co-mediating SST signaling with respect to both top-down bottom-up cortical communications and thalamocortical input processing, likely via different layer-specific cell-types. It is noteworthy that, across species, reelin is rarely co-expressed with CCK or VIP which were also investigated in this study ^92,93,96,97^.

Human reelin mRNA encodes a large, secreted glycoprotein (388 kD) ^98^ and is produced by bitufted and Martinotti GABA-ergic interneuron subtypes in lower cortical layers ^94^. In animal models, reelin enhances NMDA receptor neurotransmission through increased calcium flux ^99^, increases long-term potentiation in hippocampal slices ^100^, and its intraventricular administration to mice augments cognitive ability and dendritic spine density in hippocampus ^101^. Concomitant with compromised SST+ interneurons in layer 5 of BP, the diminished reelin expression in BP versus the SZ group in layer 5 and 6b suggests that these subjects may be differentially predisposed to circuit-specific deficits affecting experience-related synaptic plasticity in LFPC corticocortical, corticostriatal and corticothalamic circuits. While the overall understanding of layer 6b is still in its infancy, it has been implicated in mechanisms of attention and shown to have a high degree of intracortical connectivity, sending axon collaterals to layers 1 and 5 while also projecting to higher order thalamic regions ^102^. Taken together with the significant reduction in layer 4 SST mRNA of BP subjects, these findings imply the possible diagnosis-specific dysregulation of not only corticostriatal (layer 5) and long-range corticocortical circuits (layer 1), but also transthalamic circuits affecting neurocognition in BP.

Our data from BP subjects demonstrating decreased reelin expression in layer 6b versus SZ further indicates a strong diagnosis-related difference in the gating of excitatory input to pyramidal neurons projecting within the cortex and to the thalamus. Although the specific higher order thalamic region potentially innervated by LFPC layer 6b pyramidal cells is undetermined, most LPFC thalamic innervation is to the MD subregion and originates layer 6, synapsing upon thalamocortical neurons which project back to layers 4 and 5, forming cortico-thalamocortical loops ^89,103^. Such corticothalamic input optimizes working memory and improves task performance by enabling thalamocortical learning ^104^. More specifically, it has been suggested that layer 6b incorporates long-range corticocortical feedback to mediate attention and memory by regulating the selection of corticothalamocortical loops associated with decision-making^102^. These findings are also consistent with the results of our double-labeled ISH assessment showing that reductions in SST-positive cells co-expressing reelin were limited to BP with reductions in layers 1, 4, and 5 paralleling the similar deficits in total SST-expressing cells in layers 1, 4, and 5 of this group.

### CCK and VIP Expression

In control subjects, CCK-positive cells were most abundant in layers 2-4 and 6a, suggesting a heightened role in patterning long-range corticocortical transmission and corticothalamocortical loops versus corticostriatal output. VIP was more robustly expressed in upper layers and layer 4, indicating similar functional importance. The analysis of CCK mRNA expression by diagnosis determined that the single significant finding was in layer 2 of SZ subjects. These results agree with CCK mRNA reductions in some ^36,105^, but not all DLPFC non-layer-specific studies involving SZ subjects ^37^. However, when considering the reduced SST expression in layer 1 of SZ subjects in the current study, it is in line with overall greater upper versus lower cortical layer interneuron-related mRNA deficits in SZ ^106^. CCK is produced by small basket cells either co-expressing VIP or large basket cells manufacturing CCK alone with their axon terminals collectively most localized to layer 2 and “thalamic” layer 4 ^79,107–109^. Based upon our microscope-mediated visualization revealing the co-expression to be in relatively small cells, we speculate that CCK mRNA decreases reported here are not of the larger basket cell-type, but rather small basket types.

CCK+ basket cells are specialized interneurons that engage in perisomatic inhibition and, in contrast to parvalbumin+ basket cells, convey information from subcortical regions related to emotion, motivation, and autonomic states through expression of serotonin 5-HT3, CB1 cannabinoid, and nicotinic alpha7/alpha4 receptors^110^. Balancing of GABA and CCK release from these interneurons has been proposed to influence anxiety states by acting upon postsynaptic GABAA receptor subunits that mediate the anxiolytic effects of benzodiazepines and CCK2 receptors that lead to anxiogenic effects ^110^. Such diverse neurochemical drive to CCK basket cells is also partly responsible for CCK+ cell-mediated regulation of theta and gamma oscillations with their dysregulation being thought to contribute to the altered oscillatory activity within prefrontal cortex present during working memory tasks in SZ patients ^111^.

That compromised CCK+ interneurons synapse in LFPC layer 2 is of functional significance when considering the hierarchal configuration of long-range corticocortical connections proposed by the laboratory of Barbas and colleagues ^80^. According to the principles of eulaminate (highly evolved cortex) efferent targets, the most prominent LFPC projections to the anterior midcingulate (SAN hub) and pregenual cingulate/posterior cingulate (DMN hubs) largely originate in upper cortical layers and are thus potentially anatomically stronger than that compared to layer 5 where consistent expression changes were associated with BP subjects in our study. In addition, lower cortical layers of the SAN/DMN hubs constituting less evolved cortex would be expected to return robust efferents to the upper layers of the highly eulaminate LFPC. Taken together, this is conceptually aligned with the possibility that relative to bipolar illness, schizophrenia may exhibit greater CCK+ cell-mediated dysfunction of stronger reciprocal circuits associated with long-range neurocognitive function. Indeed, during resting state, LFPC functional connectivity to both the anterior midcingulate and posterior cingulate cortex is reduced in SZ patients ^112^. This provides some evidence that the magnitude of neurocognitive deficits may relate more to disrupted oscillatory activity amongst strongly interconnected network hubs and not more simply the number of local interneuron genes displaying expression changes.

Further expanding on the current single-label CCK mRNA analysis, results from our double-labeling experiments examining all CCK/VIP co-expressing cells showed significant decreases in cell density in layers 1, 2, and 6a in BP subjects. SZ subjects shared commonality with BP by displaying reduction of this cell type within layer 2 which also contained the highest degree of CCK/VIP colocalization in controls (44%). Again, based upon our microscope-mediated visualization revealing this co-expression appeared to be in relatively small cells, we speculate they are basket type. These cells likely refer to a subpopulation of CCK+ cells that synapse upon the somal regions of pyramidal cells as opposed to the disinhibitory VIP+ interneurons involved in disinhibition through contacts with SST+ cells ^41,110,113^. Consequently, this altered layer-specific interneuron population is well positioned to affect the output of long-range cortical networks and corticothalamic circuits in BP. CCK/VIP+ co-expressing subpopulations have not been well investigated in human subjects. That CCK/VIP+ cell density was highest in upper cortical layers in control subjects supports the hypothesis of CCK/VIP co-release as being particularly important for long-range corticocortical communications. Furthermore, considering our diagnosis analyses showed that most reductions in total interneurons colocalizing CCK/VIP were in BP subjects, we suggest that transcriptional regulation of key interneuron markers is not universally more notable in SZ than BP.

## Limitations

A limitation of the current study involves experimental procedures during quantification. Our method of thresholding signals above background, using consistent intensity levels across subjects, to ensure ideal cellular gene expression capture, may have resulted in quantitative loss of mRNA-positive cells having remarkably low expression signal more comparable to background intensity. Therefore, our results may be interpreted as decreased mRNA in cell populations exhibiting moderate to high gene expression. In addition, although much of our data is presented as cell density (cells/mm^2^), we cannot determine if the mRNA reductions are associated with actual neuronal loss. For example, using Nissl staining, some evidence in the anterior cingulate suggests that interneuron loss exists in SZ, especially in layers 2 and 5 ^33–35^. However, in situ hybridization analyses in DLPFC indicate that interneurons in SZ are intact, but do not express detectable levels of GAD67, SST, and PV ^39,114^. Whether reduced gene expression in LFPC of SZ and BP subjects in our analyses is associated with neuronal loss remains to be determined.

## Conclusions

The current study revealed some shared dysregulation between BP and SZ—specifically decreased SST mRNA in layer 1 in both disorders, suggesting a common top-down and bottom-up dysregulation that may affect decision-making and possible functionality of the FPN. But notably, we observed largely diagnosis-specific layer-resolved reductions in interneuron gene expression that distinguish between the two disorders.

Overall, BP subjects displayed the more appreciable gene transcription deficits, particularly in cortical layers known to regulate corticostriatal output and cortico-thalamocortical loops, with SST mRNA reduced versus controls and reelin expression was lower in BP versus SZ. In addition, density of double-labeled cells co-expressing either SST/reelin or CCK/VIP were also predominantly decreased in the BP group. These data collectively demonstrate that SZ and BP subjects have differential LFPC layer-related declines in gene expression which may be interrelated with alterations in corticocortical, corticostriatal, and transthalamic circuit functions mediating decision-making processes.

## Acknowledgements

We thank Dr. Cortney Turner and Ms. Jennifer Fitzpatrick for technical assistance on the project. This work was supported by the Pritzker Neuropsychiatric Disorders Research Consortium and the Hope for Depression Research Foundation.

## Supplementary Methods

### Fluorescence In Situ Hybridization Chain Reaction (HCR FISH)

Split-initiator DNA probes (version 2.0) were designed in our lab (based upon Choi et al., 2018; Kumar et al., 2021, Wei et al., 2023) and synthesized by Integrated DNA Technologies (Coralville, Iowa, USA). DNA hairpins conjugated with Alexa Fluor-488 (AF-488), AF-546, AF-594 and AF-647 were purchased from Molecular Instruments, Inc. (Los Angeles, CA, USA). Postmortem frozen brains were cryosectioned (30 µm thick sections) for HCR FISH. We optimized the HCR FISH method as described previously (Choi et al., 2018; Kumar et al. 2021, Wei et al. 2023). Briefly, sections were immersion fixed in 4% paraformaldehyde-saline, washed with 5x sodium chloride/sodium citrate/0.01% tween-20 (SSCTw) buffer for 3 times, 5 min each, then acetylated in 0.1M triethanolamine, pH 8.0 with 0.25% vol/vol acetic anhydride solution for 10 min. After rinsing with ddH2O, sections were de-lipidated in −20°C chilled acetone: methanol (1:1) for 5 min, washed with 5xSSCTw, and equilibrated in hybridization buffer (30% deionized formamide, 5x SSC, 9 mM citric acid (pH 6.0), 0.5 mg/ml yeast tRNA, 1x Denhardt’s solution, 10% dextran sulfate, 0.1% Tween 20) for 15 min, then incubated in hybridization buffer containing 10nM initiator-labeled probes at 37°C for 16-18 hrs. Following hybridization, sections were washed at 37°C with probe wash buffer (30% formamide, 5x SSC, 0.1% tween 20) for 3 times and twice with 5x SSCTw for 15 min each. Sections were then equilibrated in amplification buffer for 15 min (5x SSC, 10% dextran sulfate, 0.1% Tween 20). 3 µM stock fluorophore-labeled hairpins were heated at 90°C for 90 seconds and then snap-cooled to room temperature for 30 min in the dark. Snap-cooled hairpins were further diluted to 20nM final concentration in amplification buffer. Sections were incubated in amplification buffer with hairpins for 16-18 hrs at room temperature. Following the washing in 5x SSCTw twice for 30 min, sections were incubated for 5 min in TrueBlack (Biotium, Inc., Fremont, CA, USA) to quench the lipofuscin autofluorescence. Sections were then briefly rinsed in 5x SSC containing DAPI (1:10000 of 1mg/ml stock) and mounted with Vectashield antifade mounting medium (Vector Laboratories, Inc., Burlingame, CA, USA).

### Confocal Microscopy

Image stacks were acquired using an Olympus Fluoview-3000 confocal microscope. For quantitative analysis, a 10x magnification objective lens (Olympus UPLanSApo, numerical aperture (N.A.) 0.4 / working distance (W.D.) 3.1 mm) was used to acquire image-stacks (xy-dimension 1.2430 µm/pixel x 1.2430 µm/pixel and z-step of 4.0 µm; 6-7 z-slices per stack). High magnification representative images were acquired using 60x (silicone oil immersion, Olympus UPLanXApo, N.A.-1.42) objective. Image acquisition settings, mainly the PMT Voltage and laser transmissivity were selected for optimal pixel saturation to avoid excessive or weak signal and kept constant across the study.

### Image Processing and Analysis

Cell segmentation and quantification were performed across two marker panels. For the SST/Reelin analysis, tissue sections were obtained from Control (n = 17), Schizophrenia (SZ; n = 19), and bipolar disorder (BP; n = 18) subjects. For the VIP/CCK analysis, samples were derived from Control (n = 20), SZ (n = 18), and BP (n = 18) subjects.

Following HCR FISH and confocal imaging, post-processing and quantitative analysis were performed using a custom ImageJ (Fiji) (Schindelin, 2012) macro developed to segment, filter, and quantify labeled cells across defined cortical layers in postmortem human brain sections. Multichannel .tif images were generated from Olympus FV3000 .oir files via batch export. Z-projections (standard deviation) were applied to each stack, followed by background subtraction and channel splitting to isolate individual fluorescent signals (e.g., DAPI, SST, REELIN, VIP, CCK).

Each marker channel underwent Gaussian and median filtering, followed by Phansalkar local thresholding (Phansalkar, 2011) to segment punctate HCR FISH signals. Size-and circularity-based filters excluded small, irregular, or large artifact-like objects. Overlapping objects were merged with morphologically dilated small-cell candidates to refine the final segmented mask for each marker.

To ensure specificity, all segmented targets were filtered to retain only objects overlapping with segmented DAPI-positive nuclei (generated via StarDist plugin, Schmidt, 2018). Binary overlap was computed using the “Overlap Extractor” plugin in Fiji.

Colocalized populations (e.g., SST+/REELIN+ and CCK+/VIP+) were identified by logical AND operations between binary masks and further refined using size filters. Manually annotated cortical layer ROIs (drawn based on DAPI cytoarchitecture) were batch-imported per section. Cell counts for each marker and colocalized population were quantified per layer using Analyze Particles, and cortical layer areas were measured to compute normalized cell densities. All results, including raw counts, area measurements, and segmented masks, were exported for downstream statistical analysis.

The full analysis pipeline, including the main Analysis macro, and ROI drawing/saving macro, is available at: https://github.com/vivekk-MNI/Postmortem_Human_Brain_HCR_FISH_Cell-_Quantification_Macro_Fiji-ImageJ.git

---

**Table 1.** Patient demographics and tissue storage time associated with diagnosis. . Categories for gene expression of SST and reelin in CTR, SZ, and BP are listed in the upper row while that for CCK and VIP appear in the bottom row. AA; African American, C; Caucasian, CCK; cholecystokinin, CTR; control, BP; bipolar disorder, F; female, M; male, H; Hispanic, N; number of subjects, PI; Pacific Islander, SZ; schizophrenia, SST; somatostatin, VIP; vasoactive intestinal peptide.

|  | Diagnosis | N | M/F | Race | Age | pH | PMI | Storage (years) |
|---|---|---|---|---|---|---|---|---|
| SST/Reelin | CTR | 17 | 17/0 | (15)C,(1)AA,(1)H | 50.59±8.16 | 6.85±0.13 | 24.39±4.69 | 18.25±5.17 |
|  | SZ | 19 | 19/0 | (16)C,(1)H,(1)A, (1)PI | 47.16±14.04 | 6.86±0.16 | 20.29±8.87 | 19.26±3.57 |
|  | BP | 18 | 13/5 | (18)C | 46.33±13.02 | 6.76±0.16 | 22.87±7.39 | 21.28±4.38 |
| CCK/VIP | CTR | 20 | 20/0 | (18)C,(1)AA,(1)H | 50.95±6.98 | 6.81±0.14 | 23.07±5.37 | 18.80±5.05 |
|  | SZ | 18 | 18/0 | (15)C,(1)AA,(1)H,(1)PI | 48.05±14.08 | 6.87±0.16 | 20.46±9.09 | 19.28±3.68 |
|  | BP | 18 | 15/3 | (18)C | 44.33±12.88 | 6.78±0.16 | 23.09±7.22 | 20.78±4.52 |

**Supplemental Fig. 1.**
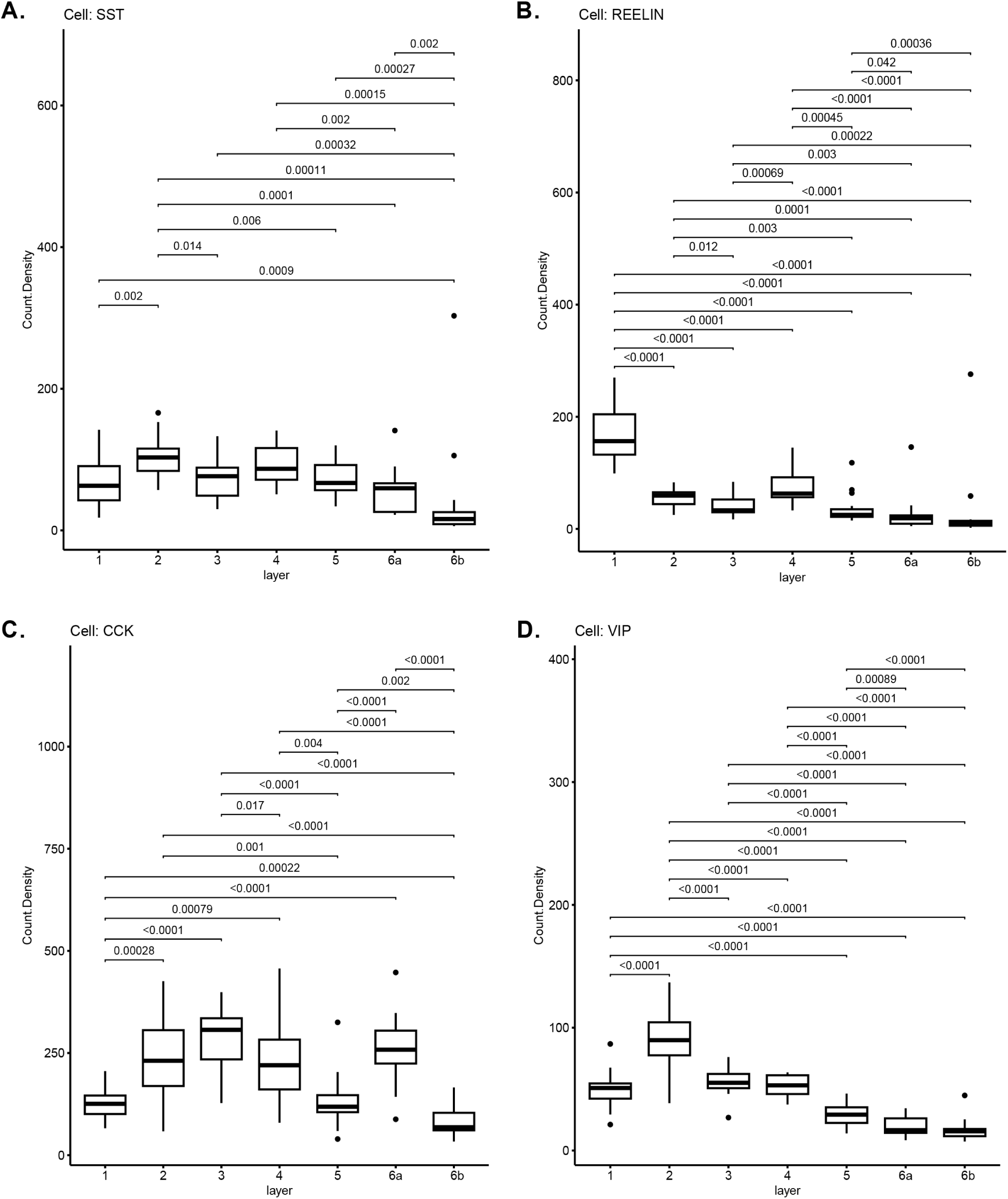
The density of LFPC cells expressing mRNA for either SST, reelin, CCK, or VIP by cortical layer in control subjects. Box plots show the median number of positively labeled cells for gene expression with the upper 75% and lower 25% quartiles at the top and bottom of the box, respectively. Dots represent outliers defined as 2x standard deviation above 1.5x the quartile value. **A.** The cellular density of cells expressing SST in cortical layer 2 was significantly higher than all other layers, except layer 4. Layer 6b exhibited the lowest median cell count which was significantly lower compared to all other layers. **B.** Density of reelin-positive cells in cortical layer 1 was larger than all other layers. Layer 4 cell density was increased compared to layers 3 and lower layers 5, 6a, and 6b. **C.** CCK-positive cells in cortical layers 2, 3, 4, and 6a were significantly more abundant compared to layers 1, 5, and 6b. **D.** Cells expressing VIP were significantly higher in layer 2 compared to all other layers. VIP-positive cells positioned in layers 1, 3, and 4 were more densely expressed than layers 5, 6a, and 6b. CCK; cholecystokinin, LFPC; lateral frontal pole cortex, SST; somatostatin; VIP; vasoactive intestinal peptide.

**Supplemental Fig. 2.**
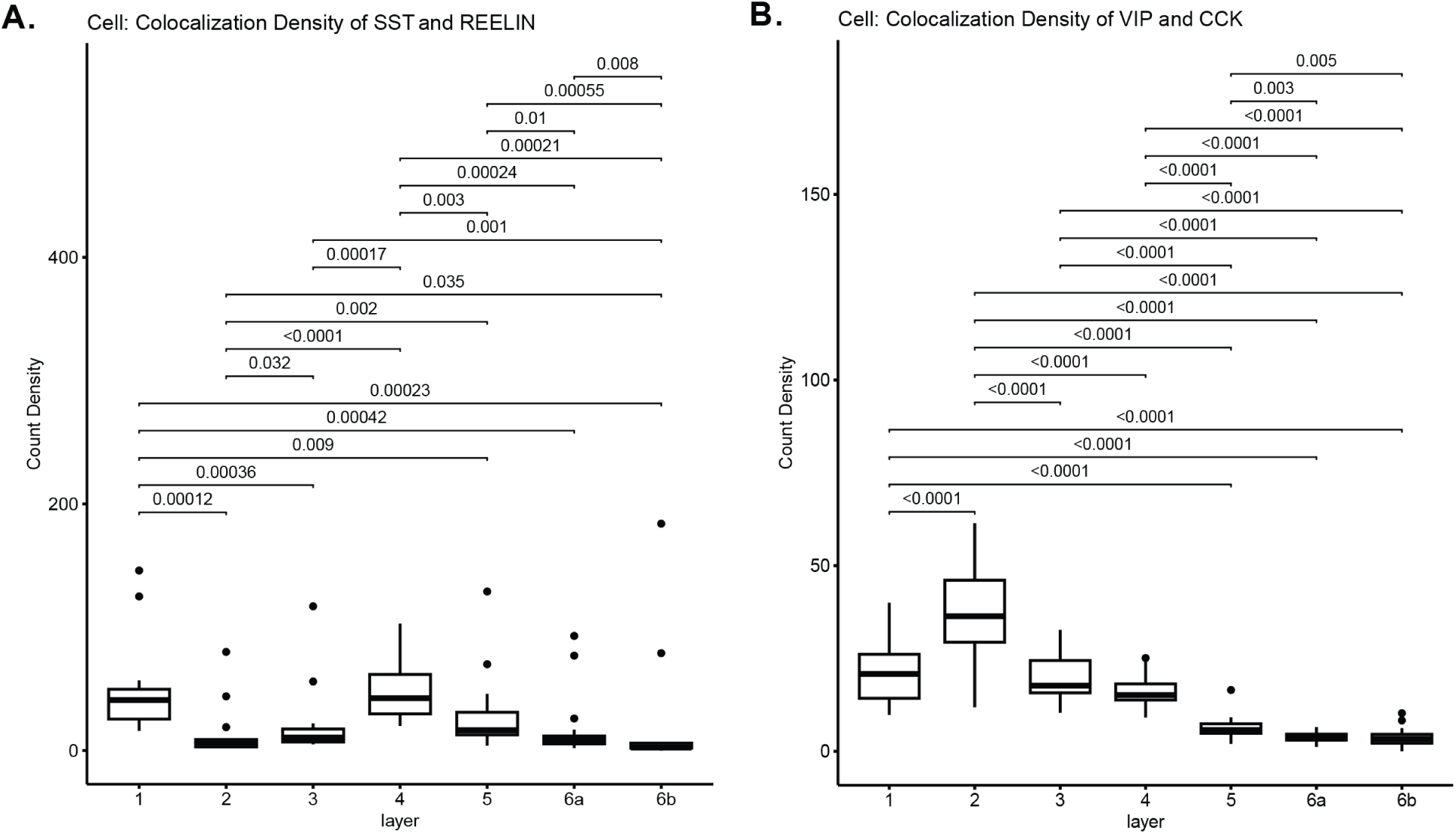
The density of LPFC cells co-expressing either SST and reelin or CCK and VIP by cortical layer in control subjects. Box plots show the median number of positively labeled cells for gene expression with the upper 75% and lower 25% quartiles at the top and bottom of the box, respectively. Dots represent outliers defined as 2x standard deviation above 1.5x the quartile value. **A.** The cellular density of SST colocalization with reelin in cortical layers 1 and 4 was significantly higher than all other layers while that in layer 4 showed the highest median, compared to layer 1, where cell count was significantly greater than remaining layers 2, 3, 5, 6a, and 6b. **B.** CCK-positive cells co-expressing VIP were most abundant in cortical layer 2 where they were more dense than all other layers. This cell-type was significantly denser in layers 1 and 3 compared to 5, 6a, and 6b. CCK; cholecystokinin, LFPC; lateral frontal pole cortex, SST; somatostatin; VIP; vasoactive intestinal peptide.

**Supplemental Fig. 3.**
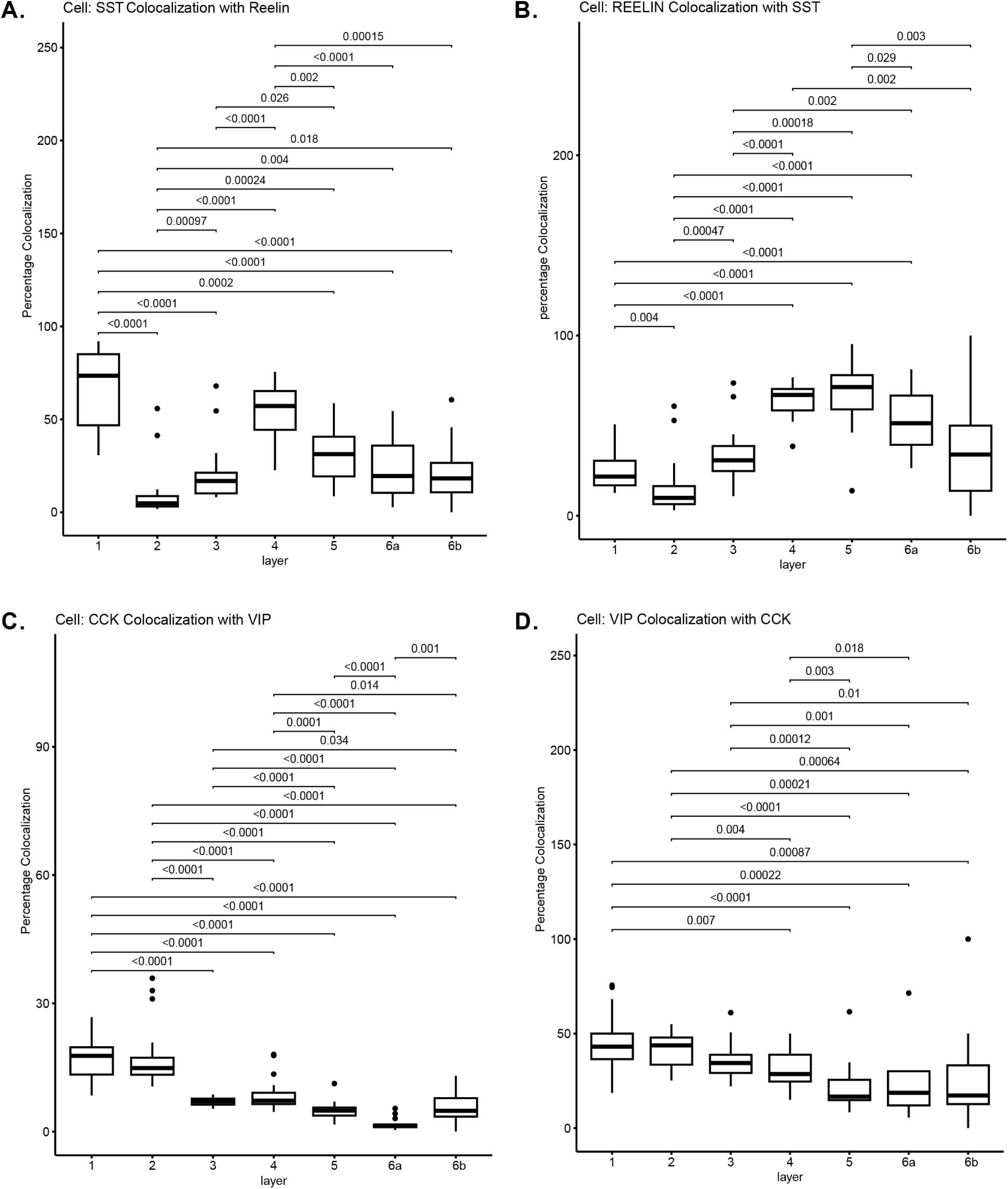
The relative percentage of LPFC cells co-expressing either SST and reelin or CCK and VIP by cortical layer in control subjects. Box plots show the median number of positively labeled cells for gene expression with the upper 75% and lower 25% quartiles at the top and bottom of the box, respectively. Dots represent outliers defined as 2x standard deviation above 1.5x the quartile value. **A.** The percentage of cells showing SST colocalization with reelin in cortical layers 1 and 4 was significantly higher than all other layers while that in layer 1 showed the highest median, compared to layer 4. **B.** The reverse analysis showed the percentage of reelin-positive cells colocalizing with SST was significantly higher in layers 4 and 5 compared to all other layers, except 6a. Layers 4 through 6a generally exhibiting heightened percent colocalization versus upper layers 1 through 3. **C.** The percentage of CCK-positive cells that exhibited co-expression VIP demonstrated that layers 1 and 2 were significantly higher than all other layers. **D.** Assessment of VIP-positive cells co-expressing CCK showed that the greatest percentage of these cell-types was found in the upper layers 1 and 2 demonstrating significant differences versus layers 4 through 6b. CCK; cholecystokinin, LFPC; lateral frontal pole cortex, SST; somatostatin; VIP; vasoactive intestinal peptide.

**Supplemental Fig. 4.**
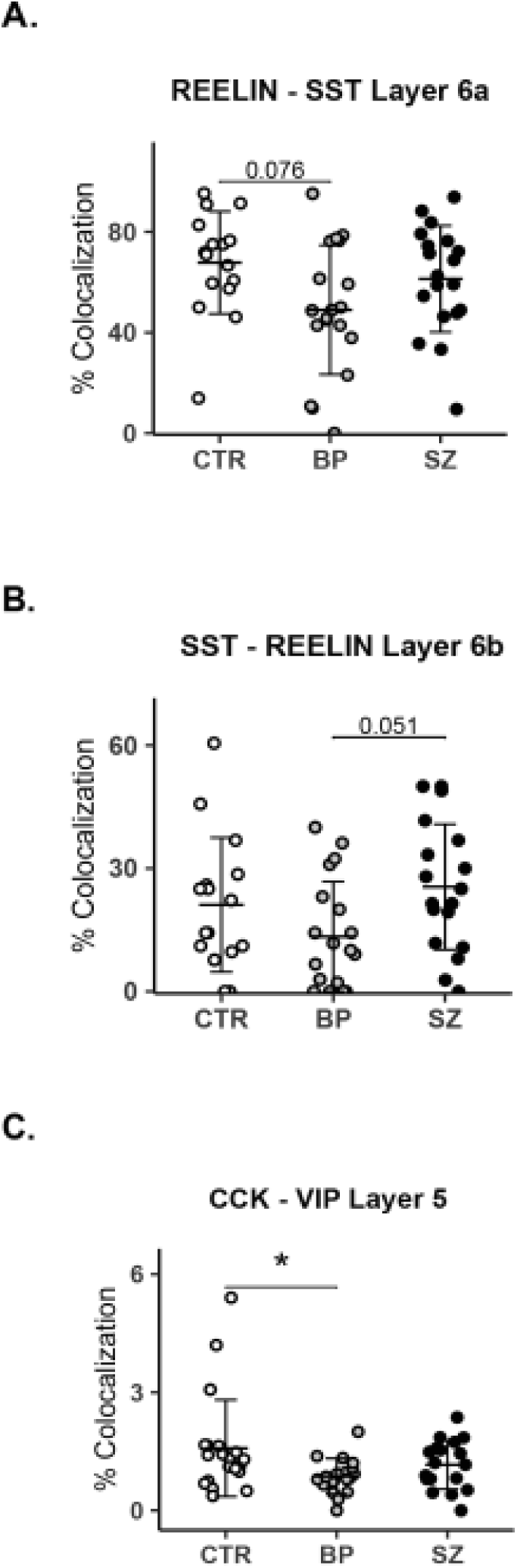
Percentage of cortical layer-based colocalized cells in the LFPC of psychiatric subjects versus controls. **A.** BP subjects demonstrated a non-significant trend for decreased percentage of reelin cells colocalized with SST in layer 6a, p=0.076. **B.** There was a strong trend for reduced percentage of SST cells co-expressing reelin in BP subjects versus SZ, p=0.051. **C.** The percentage of CCK cells colocalized with VIP was significantly lower in layer 6a of BP subjects versus controls, p=0.015. BP; bipolar disorder, CTR; control, CCK; cholecystokinin, LFPC; lateral frontal pole cortex, SZ; schizophrenia, SST; somatostatin; VIP; vasoactive intestinal peptide. Asterisk represents p=0.015.

